# The influence of startReact on long-latency reflexive muscle activation during the transition from posture to movement

**DOI:** 10.1101/180554

**Authors:** Claire F. Honeycutt, Vengateswaran J. Ravichandran, Eric J. Perreault

## Abstract

Many motor tasks involve transitions from posture to movement such as shifting from holding an object to setting it on a surface. Although many movements are voluntary processes, they are facilitated through motor pathways that span the continuum between reflexive and voluntary control. The mechanisms driving the long-latency stretch reflex (LLSR) have remained hotly contested. Recently, startReact has been shown to influence the fast muscular response to perturbations (Ravichandran et. al. 2013). The objective of this study was to evaluate how the LLSR and startReact impact the muscular response during the transition from posture to movement. We hypothesized that both the LLSR and startReact would be involved throughout the transition from posture to movement; however there would be a clear transition from LLSR dominance during posture to startReact dominance near/during movement. We tested this hypothesis using perturbations of elbow posture at various times before a fast ballistic extension movement. We found clear evidence that both the LLSR and startReact components were influenced changes in the late long-latency time window. The results provide insights on how the nervous system regulates involuntary responses to perturbations during the transition from maintenance of arm posture to movement.

**New and Noteworthy:** We recently demonstrated that the startReact effect, the startled release of a planned movement, can influence the muscular response during the LLSR. Here we observe how the influence of startReact can change from the transition from posture (no/little influence) to movement (strong influence). While not all paradigms trigger a startReact effect, this work demonstrates that when present startRect can have a profound effect on the overall muscle activity - even obscuring the traditional LLSR response.

## Introduction

A wealth of motor tasks involve transitions from posture to movement. One example is that which occurs when an arm is used to support an object against gravity, prior to placing it on a shelf. During this transition, the primary objective of the arm changes from the control of posture and stability to the control of movement. Although many movements are voluntary processes, their control is facilitated through numerous motor pathways that span the continuum between reflexive and voluntary control.

Human reflex studies have focused extensively on limb-stability tasks (i.e. those in which limb stability is compromised; also referred to as goal-dependent tasks) (Lee and Tatton, 1975, Doemges and Rack, 1992; Dietz et al., 1994; Perreault et al., 2008; Krutky et al., 2009; Pruszynski et al., 2011; Nashed et al., 2014; Weiler et al., 2016) and instruction-dependent tasks (i.e. those in which the subject is instructed how to react to a perturbation) (Hammond, 1956; Crago et al., 1976; Colebatch et al., 1979; Newell and Houk, 1983; Koshland and Hasan, 2000; MacKinnon et al., 2000; Lewis et al., 2006; Mutha et al., 2008; Pruszynski et al., 2008; Pruszynski et al., 2009; Shemmell et al., 2009). During both limb-stabilizing and instruction-dependent tasks, the long latency stretch reflex (LLSR) is modified in a context-appropriate manner. In limb-stabilizing tasks, reflex excitability increases to enhance the stiffness and stability of the arm; during instruction-dependent tasks, the perturbations evoke a motor response in the long latency time period that facilitates the motor plan.

The mechanisms driving the distinct responses during limb-stabilizing and instruction-dependent tasks remain hotly contested. This is partly driven by the existence of two conflicting hypotheses about the dominant contributor to LLSR modulation. The first hypothesis is that LLSR modulation is driven by a pathway that contributes to the regulation of limb mechanics such as compensating for inter-joint coupling or interactions with challenging task environments (Kimura et al. 2006; Prochazka et al. 2000; Pruszynski et al. 2011; Shemmell et al. 2009). This component has known cortical origins (Pruszynski et al. 2011; Shemmell et al. 2009) and has a strong and dominant presence during limb-stabilizing tasks.

The second hypothesis is that LLSR modulation is driven by triggered reactions (Crago et al. 1976; Ravichandran et al. 2013). The triggered reaction hypothesis has been discounted in large part because triggered reactions are only present during certain tasks e.g. those where the subject is preparing movement. We have recently posed an explanation for this. When a subject is in a state of motor planning, a startling stimulus (e.g. perturbation of the arm, loud acoustic stimulus) produces an early release of planned movement. These startle-evoked-movements, referred to as startReact, only occur in a state of motor planning and have been evaluated extensively in the literature (Valls-Solé et al., 1999; Siegmund et al., 2001; Carlsen et al., 2004; Honeyuctt et al. 2012; Honeycutt et al. 2013; for review, see Valls-Solé et al., 2008). It is important to note that startReact is not a “stretch reflex” but rather an early release of planned movement that coincides with the LLSR. As the term triggered reactions has been associated as being a component of the LLSR in the literature (and not a separate phenomenon), we will use the term startReact in this article to refer to the startle-evoked-movement phenomenon discovered by Valls-Sole.

Until recently, the limb-stabilizing modulation of the LLSR was perceived as the only neural response following a stretch perturbation and triggered reactions (or startReact) were seen as an artifact rather than an actual phenomenon. This was mainly due to the fact that there was no way to identify the presence or absence of startReact. This was further compounded by two factors: (1) the observation of startReact only occurs in the presence of a movement plan and (2) the use of average responses over a number of stretch reflex trials that contained both startReact and limb-stabilizing LLSRs instead of visualizing these as two separate phenomenon.

Our recent work poses a solution. StartReact is accompanied by activity in the sternocleidomastoid (SCM) muscle. Therefore we can identify the situations where startReact is present to influence the response. We recently utilized this methodology to demonstrate that the instruction dependent modulation during the long-latency stretch reflex time window is associated with indicators of startle (Ravichandran et. al. 2013). For example, SCM activity is not present when displacement perturbations are applied during the maintenance of limb posture (Ravichandran et. al. 2013) indicating that startReact is not influential during these tasks. It is important to note that while we can determine when startReact is present or absent, the presence of startReact does not preclude additional influence from other mechanisms. In conclusion, monitoring SCM activity provides a means to identify when startReact is present and therefore influencing EMG activity during the LLSR.

The objective of this study was to evaluate how the LLSR and startReact are impacted during the transition from posture to movement. Previous works demonstrated that the amplitude of the LLSR is modulated by the time of perturbation during the transition phase (Mortimer et al. 1981) and by the intended direction of movement (Bonnet 1983). Though these studies found increased LLSRs preceding volitional movement, they could not account for the relative contributions of startReact. Since the postural demands remain constant prior to movement (Brooke et al. 2000), we hypothesized that both the LLSR and startReact would be involved throughout the transition from posture to movement; however there would be a clear transition from LLSR dominance during posture to startReact dominance near/during movement. We tested this hypothesis using perturbations of elbow posture at various times before a fast ballistic extension movement. The results provide insights on how the nervous system regulates involuntary responses to perturbations during the transition from maintenance of arm posture to movement. As the LLSR and startReact have unique neural mechanisms, understanding how each is regulated may inform our understanding about how movement transitions are affected during neurological disease and injury.

## Methods

### Ethics approval

Experiments were performed on eight right-handed, able-bodied subjects (age: 25 ± 3, 4 females and 4 males) with no known neurological disorders. The right arm was used for testing for all subjects. All protocols were approved by the Northwestern University Institutional Review Board (IRB Protocol STU00009204) and required informed written consent.

### Apparatus Setup

Subjects were seated comfortably with the trunk secured to an adjustable chair (Biodex, NY) using padded straps such that their initial posture was at 70° shoulder abduction, 0° shoulder flexion, and the elbow was at 90° flexion. The wrist was immobilized in a neutral position using a rigid custom-made plastic cast (Fig. 1). The cast was directly attached to a force sensor (45E15A4-I63-AF 630N80; JR3 Inc, Woodland, CA) mounted on a rotary motor (BSM90N-3150, Baldor Electric Company, WV), aligned such that the motor axis was in line with the elbow flexion/extension axis. The rotary motor was attached to a 10:1 gear head (AD140-010-PO/BaldorBSM90N3; Apex Dynamics, Taiwan ROC). This allowed the encoder to record position with a resolution of 3.6 x 10^-3^ °.

**Figure 1:**
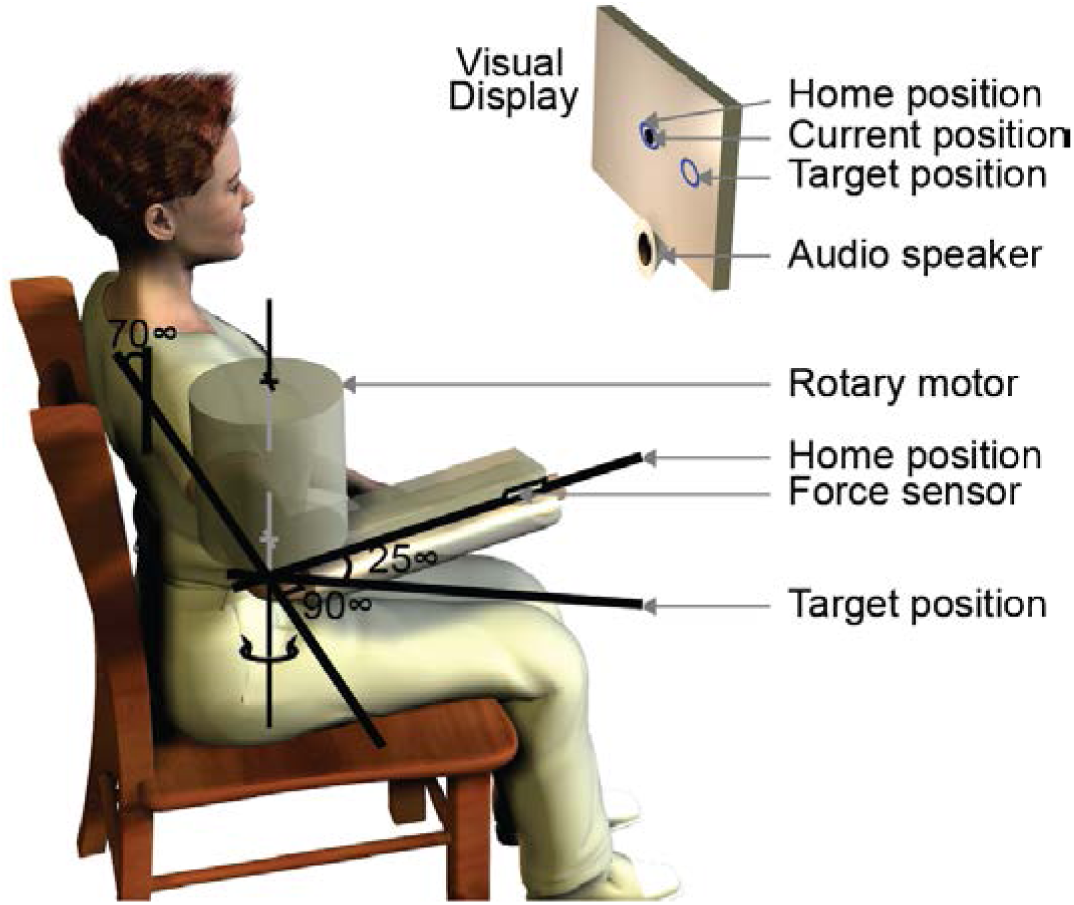
The setup used for the experiment. The shoulder straps and the lap belt used for restraining the subject are not shown in the figure. The shoulder horizontal flexion was at 0° and the wrist was slightly pronated. The rest of the joint angles are shown in the figure.

The rotary motor was configured as an admittance servo, consisting of an inner position control loop with a high stiffness (> 30 kNm/rad) and torque feedback that allowed compliant environments to be rendered. This combination allowed us to superimpose precise position perturbations on top of any ongoing motion. A compliant configuration with a rendered stiffness of 0 Nm/rad, a moment of inertia was set to 0.2 kgm^2^/rad, and a critical damping was used to allow subjects to attain the initial target and to make free reaching movements. The compliant configuration was used during all times (initial posture setting and reaching movement). The rigid configuration was used during the application of perturbations such that the identical perturbation could be applied to each subject (i.e. the motor was stronger than the subject so each subject was exposed to the same perturbation regardless of strength and their reaction time). The effective stiffness of the superimposed perturbations was the same as the inner position control loop. Physical stops limited the actuator to 20° of flexion and 45° of extension from the nominal position. Software limits were implemented to prevent motion 10° before contact with the physical limits. To ensure subject safety, both the subject and the experimenter were provided with their own respective stop buttons that cut power to the motor.

Surface electromyographic (EMG) activity was recorded from the triceps lateral head, and the left sternocleidomastoid (SCM) using bipolar Ag/AgCl electrodes (Noraxon Dual Electrodes, #272, Noraxon USA Inc., AZ). EMGs were amplified and conditioned using a Bortec AMT-8 (Bortec Biomedical Ltd., Canada), with a band-pass filter of 10-1,000 Hz. The resulting signals were anti-alias filtered using 5^th^ order Bessel filters with a 500 Hz cut-off frequency and sampled at 2500 Hz using a PCI-DAS1602/16 (Measurement Computing, MA).

Visual feedback of the current elbow angle, the starting position (90° elbow flexion) and the target position (65° elbow flexion) was provided via a computer monitor placed 25 cm in front of the subject (Fig.1).

Auditory signals were used to instruct the subject to prepare to move (*WARNING* signal) and to make the movement (*GO* signal). Both WARNING and GO signals were presented via Sonalert SC628ND speakers (Mallory Sonalert Products Inc., IN) placed 25 cm in front of the subject. The peak intensities of WARNING, and GO signals near the subject’s ears were approximately 80 dB. The intensities were measured using a digital sound level meter (Model 407730, Extech Instruments Corp, MA). The duration of all auditory signals was limited to 40 ms.

### Protocols

Our experiments were designed to quantify how the nervous system regulates the LLSR and startReact during the transition from maintaining a set arm posture to arm movement. This was accomplished by applying elbow perturbations during various time points in the transition period from postural maintenance to the onset of ballistic elbow extension movements.

The experiment consisted of a *movement-training* phase and a *movement-testing* phase. In all phases the rotary motor was set to simulate a compliant environment (stiffness = 0 Nm/rad and critically damped), and subjects held a posture or produced a movement against a load that required them to exert a constant extension torque of 2 Nm throughout the experiment.

In the movement-training phase, subjects performed 40 ballistic movements from their initial position (Home position, Fig. 1) to a 25° extension target (Target position, Fig.1). The WARNING signal (80 dB tone) was provided only after the subjects had held the Home position ± 1° for a random duration between 0.5-1 s. After a randomized time interval of 2.5-3.5 s following the WARNING signal, a GO signal (80 dB tone) was provided, instructing the subject to move to the target as fast as possible. Subjects were also told that the reaction times were more crucial than the accuracy of the reach, but were instructed explicitly to avoid undershooting the target and received appropriate feedback (e.g. told to move faster or more accurately) to assist in that task.

The movement-testing phase was identical to the training phase with the exception that ramp-and-hold elbow flexion perturbations were applied in 25% of non-consecutive trials, distributed randomly. Perturbations were applied at one of 5 possible time points (Fig. 2): 500 ms before WARNING, 500 ms after WARNING, 500 ms before GO, 100 ms before GO, and at GO. All perturbations had a displacement of 6°, a velocity of 60°/s and a hold time of 250 ms after the end of the ramp.

**Figure 2:**
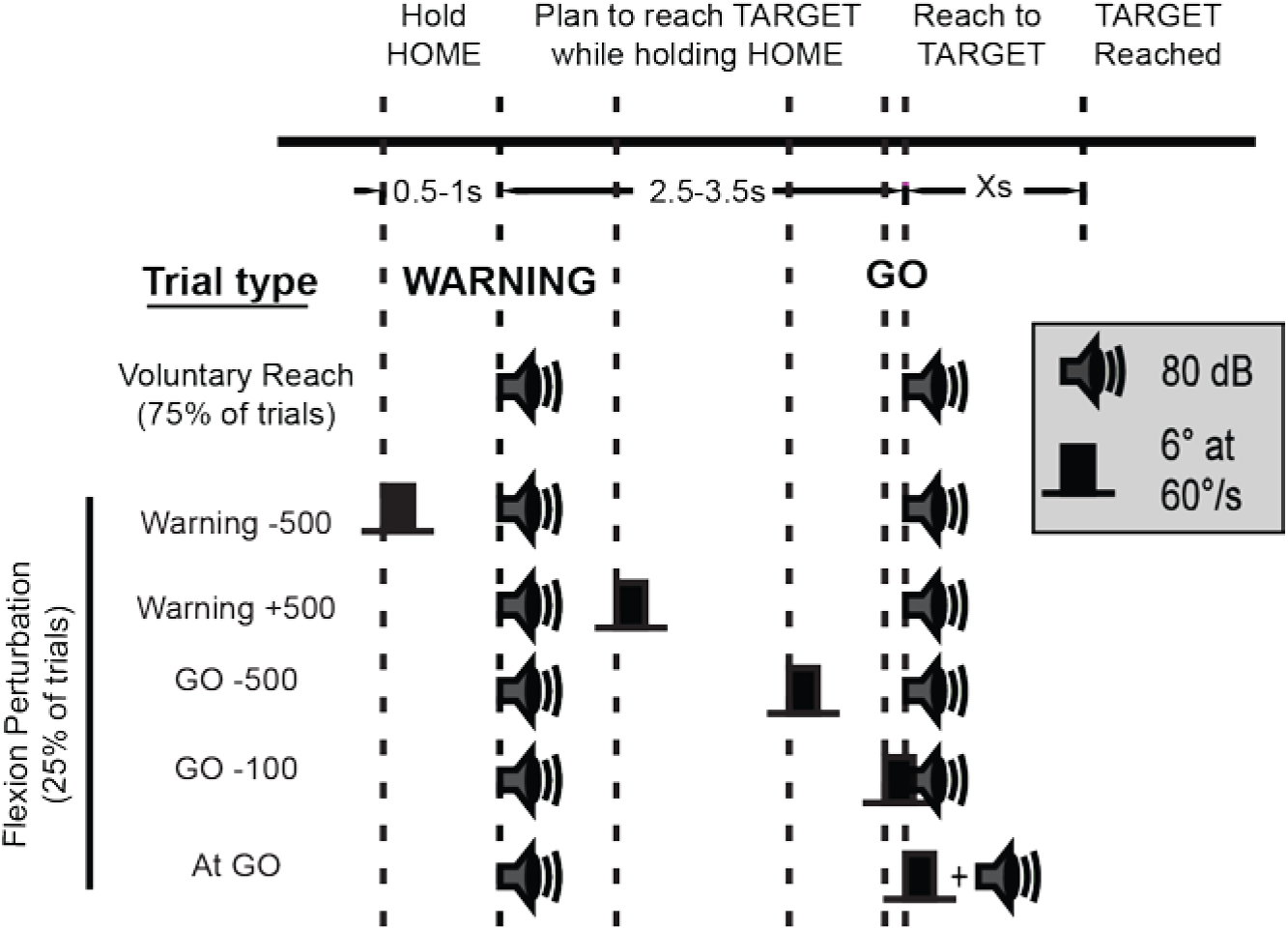
Schematic timeline of the experiment. The different types of trials used in the experiment are depicted. During voluntary reach trials, subjects got into the Home position, waited for the WARNING signal, then prepared to reach to the Target position as soon as they received the GO signal. In perturbation trials, subjects were also given a flexion perturbation in one of the 5 listed time-points, relative to the onset of WARNING/GO signals.

To minimize fatigue, subjects were limited to a total of 200 reaching trials, split into 5 blocks of 40 trials. A minimum of a 1-minute break was enforced between blocks. Subjects also had the option of resting at any point in the experiment.

### Data Analysis

For each trial, the mean values of the EMG signals were removed and the resulting signals rectified. We calculated the mean amplitude of the rectified EMG in the triceps brachii over the following time intervals. The period 100 ms before the onset of perturbation was used to quantify the background muscle activity. The short latency reflex amplitude was quantified from 20-50ms while the long-latency was split into two periods: early (50-75) and late (75-100ms). It has been previously reported that startReact is active predominately during the late time window (Ravichandran et al. 2013). Therefore, separation into two windows allows us to distinguish between startReact and LLSR contributions. Trials with background activity above or below two standard deviations from the average background for that subject were excluded from further analysis. This was used to detect early movement onsets prior to the GO cue. This process eliminated fewer than 10% of the trials in all subjects.

For each trial, we calculated onsets of the EMG response in the triceps and the sternocleidomastoid. EMG onsets were defined as the point at which the EMG activity rose above the background activity. For each individual trial, EMGs were plotted along with a line representing the background activity and another representing two standard deviations from background. The point at which the activity first exceeded the two standard deviations was first noted and visually adjusted back to the first point at which the activity deviated from the baseline.

Activity in the sternocleidomastoid was used to identify the presence of startReact. This was accomplished by separating all perturbation trials into those with activity in the sternocleidomastoid (*SCM+* trials) and those without (*SCM-* trials). Trials were categorized as SCM+ only if the onset of activity in the sternocleidomastoid was earlier than 120 ms after perturbation onset. A few (<2%) of the voluntary reach trials were classified SCM+ and not included for further analysis. As both LLSR and startReact are active during the late long latency window, we compare responses for SCM+ and SCM-thereby allowing us to distinguish between the effects of startReact and LLSR.

### Statistical Analysis

Our general hypothesis was that both the LLSR and startReact would be involved throughout the transition from posture to movement; however there would be a clear transition from LLSR dominance during posture to startReact dominance near/during movement. We specifically tested this hypothesis by evaluating amplitude changes from different time periods. We evaluated amplitude changes during different time-before-GO conditions evaluating the “null hypothesis” that no change had occurred. This was accomplished using a generalized linear mixed effects model to determine significance of the changes in the triceps EMG amplitude and probability of eliciting an SCM response. We created a generalized linear mixed effects model with time of onset of perturbation (time-before-GO) and the presence/absence of sternocleidomastoid activity (SCM) as a fixed factors and the subject as a random factor. In accordance with recent standardization of statistical practices, all individual trials were included in our analysis of the linear mixed-effects models (Hedeker and Gibbons 2006). This method has been shown to be more rigorous and powerful than using a single mean for each subject. The use of all trials allows more independent information than a single measurement decreasing the probability of statistical error by capturing all the variability within a data set. Additionally, the mixed-effects models take into account the number of trials into the analysis ensuring that data are not misrepresented or inflated due to differences in trial number across subjects (due to differences in SCM+/-classifications) in unbalanced data sets. Most relevant to this paper, the startle-related component is not present during all trials and therefore it is important to differentiate which trials where startle is and is not present in order to accomplish the objectives of this paper.

All four amplitude measures (background, short latency, early long-latency, and late long-latency)(Figure 3), the probability of SCM+ (Figure 6), and amplitudes of SCM+/-trials (Figure 7) were tested for significance as the dependent variables. Statistical significance was tested against a p-value of 0.05. Tukey post-hoc tests were used to obtain the contrast features of the ANOVA. All of the statistical analyses were performed using R (R Development Core Team, 2006). All error bars represent standard error based on total number of trials from all participants.

## Results

### Long-latency responses increase as movement time approaches

The flexion perturbations elicited reflex responses in the triceps brachii muscle of all subjects, but the amplitude of these responses varied depending on when they were applied relative to the GO signal (time-before-GO), particularly during the late long-latency response (Fig. 3). The background activity remained constant across all time-before-GO conditions (F_4,354_ = 0.53; p= 0.71). The short latency response amplitude did vary across time-before-GO conditions (F _4,354_ = 4.3; p = 0.002). However, post-hoc comparisons revealed that only the response elicited synchronous with the GO cue was different from the three earlier perturbation times [Warning – 500 (Δ = 0.02 mV, p = <0.001), Go – 500 (Δ = 0.01 mV, p = 0.03), and Go – 100 (Δ = 0.01, p = 0.03)]. All other comparisons did not reach significance (max Δ = 0.009, all p > 0.46). The amplitude of the early long-latency response was also sensitive to the time at which the perturbation was delivered (F _4,354_ = 5.4; p = 0.0003). Specifically, perturbations delivered at the two earliest time points (Warning – 500 and Warning + 500) were significantly different from those delivered closer to the onset of the planned movement (Go – 100 and At GO) (max Δ = 0.03 mV, all p < 0.03). The late long-latency amplitude was the most sensitive to when the perturbation was applied (F _4,354_ = 29.27; p < 0.0001). There were significant differences in the response elicited by all perturbation times (max Δ = 0.14 mV, p < 0.03) except the two earliest (Warning – 500 and Warning + 500; Δ = 0.005 mV, p = 0.99) and the two latest conditions (Go – 100 and At GO; Δ = 0.01 mV, p = 0.98). These results are consistent with the findings from previous studies that show increasing amplitude of the LLSR prior to movement onset (Gottlieb and Agarwal 1980; Mortimer et al. 1981). Since the background was not changing in our experiment, we can rule out the possibility that this increase in long-latency response comes from a change in background activity (Hallett et al. 1981).

**Figure 3:**
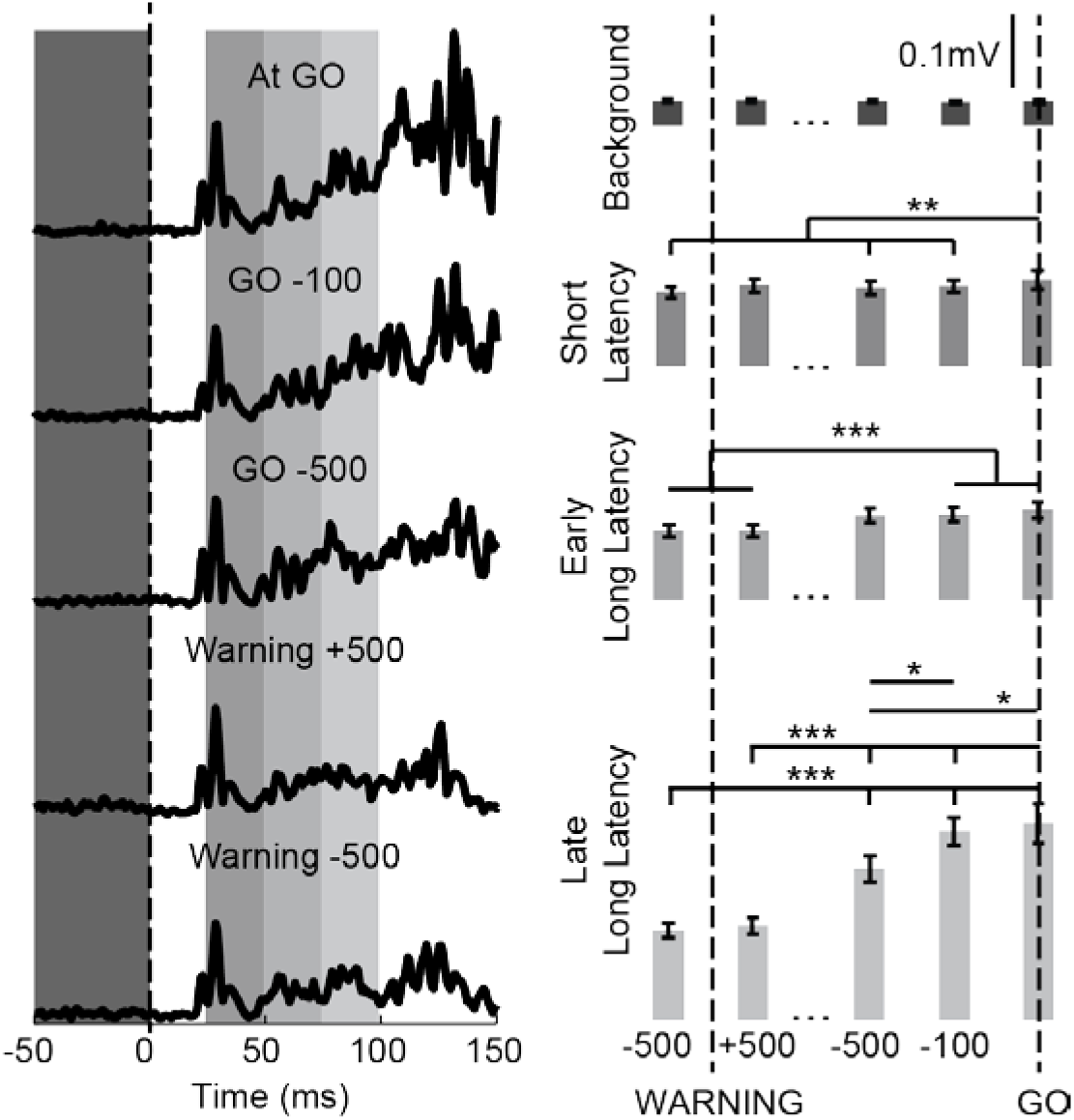
Sample data from one subject and group data showing the triceps response to flexion perturbation applied at 5 different points of time. On the left is representative EMG data from one subject. Each trace is the average of between 7 and 10 responses for each of the perturbation times relative to the GO cue. The onset of perturbation is at 0 (dashed vertical line). The right panel shows the average amplitudes of between 52 and 80 trials (mean ± standard error) across all subjects from each of the analysis windows; the X-axis represents time of perturbation onset.

### Perturbations evoked sternocleidomastoid muscle activity

Perturbations delivered closer to the GO cue evoked more consistent responses in the SCM muscle. We categorized each perturbation trial as SCM+ or SCM-, based on the presence or absence of activity in this muscle (Fig. 4). Across all subjects, the probability of eliciting EMG activity in the sternocleidomastoid increased as perturbations were delivered closer to the GO cue (Fig. 5; F_4,28_ = 53.8; p < 0.0001). The probabilities of eliciting SCM activity were significantly different across all perturbation time points (all p < 0.0001) except the two earliest conditions (Warning – 500 and Warning + 500; Δ = 0.03 mV, p = 0.99) and the middle two conditions (GO – 500 and GO – 100; Δ = 0.11 mV, p = 0.37). During maintenance of arm posture (500 ms before WARNING), when there was no movement plan, the probability of evoking a startle was zero in all subjects. However, after the WARNING signal, i.e., once a reach target was provided, the probability of evoking a startle increased gradually until it peaked at an average of 0.78 when the perturbation was applied at GO.

**Figure 4:**
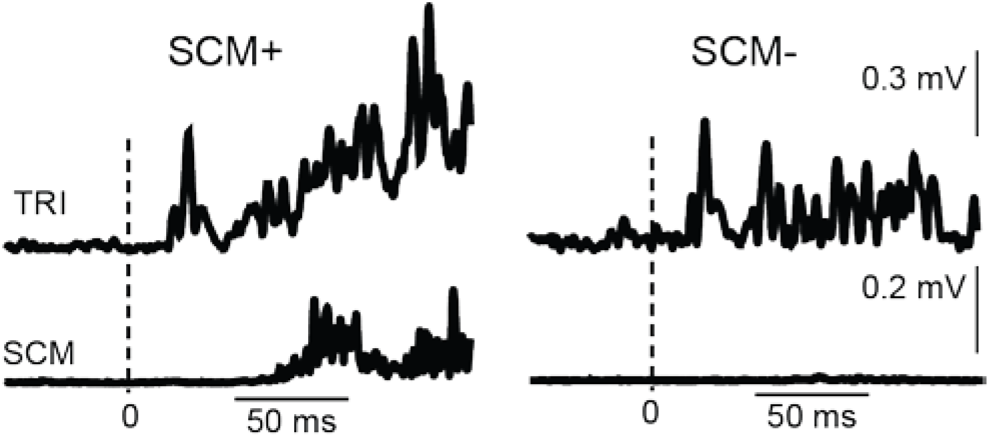
Averaged EMG response of triceps brachii and sternocleidomastoid muscles to perturbation applied 500 ms prior to GO. The onset of the perturbation is at 0. The top traces shows the average EMG response in the triceps muscle to a flexion perturbation during SCM+ (left) and SCM-(right) trials. The bottom EMG traces show the average sternocleidomastoid muscle activity during SCM+ (left) and SCM-(right) trials.

**Figure 5:**
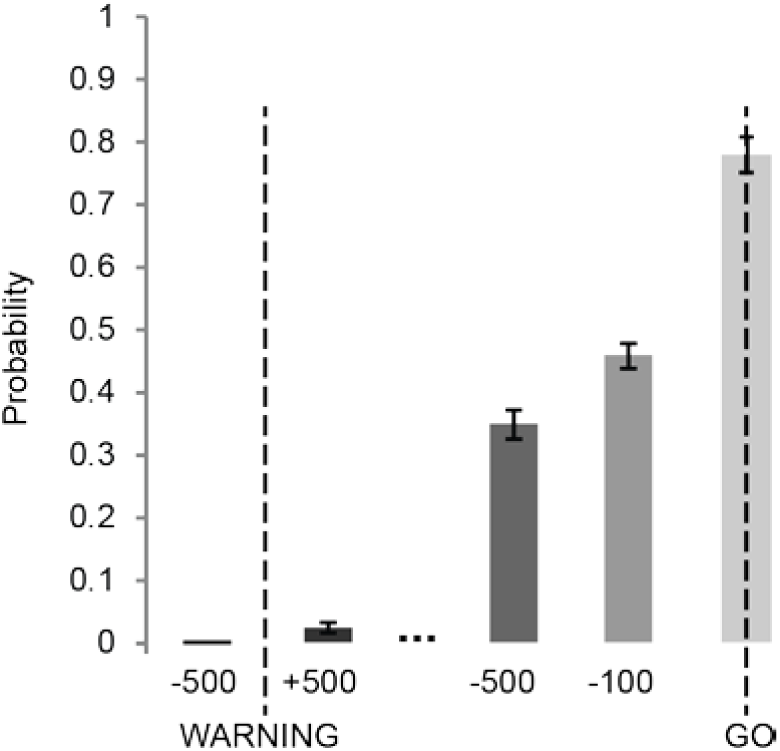
Probability of evoking a response in the sternocleidomastoid muscle. Bars indicate the mean ± standard error of the probabilities across all subjects. Separate results are plotted for each of the 5 different times that perturbations were applied during the preparation for movement.

### Influence of startle-mediated pathways on the long-latency stretch-perturbation response

The perturbation-evoked activity in the triceps was larger during the SCM+ trials than in the SCM-trials, an effect that was largest during the late long-latency window. SCM+ trials were larger during both the early long-latency (Δ = 0.04 mV; F 1,357 = 49.7; p < 0.0001) and late long-latency (Δ = 0.13 mV; F _1,357_ = 123.4; p < 0.0001) windows though the difference was 3.25 times larger in the late long-latency time window compared to the difference in the early long-latency response. These results further support previous work indicating that the startle-related components (SCM+ trials) make their most significant contribution during the late long latency response (Ravichandran et al. 2013). Therefore, the impact of SCM+ was analyzed in further detail during the late long-latency window.

The amplitude of the triceps EMG response during the late long-latency time period depended on both the activity in the sternocleidomastoid muscle as well as the time at which the perturbation was applied relative to the onset of the GO cue (sample data: Fig.6; group results: Fig 7). There were no SCM+ trials evoked during the WARNING – 500 condition and only 2 SCM+ trials in one subject during the WARNING + 500 condition. Therefore, these conditions were excluded from further analysis. Within the remaining three conditions (GO – 500, GO – 100, At GO), there were influences of both condition (F_2,197_ = 6.65; p = 0.0016) and startle (F_1,197_ = 28.4; p<0.0001) but no interaction (F_2,197_ = 1.23; p = 0.29). All SCM+ trials showed larger amplitudes compared to SCM-trials. In SCM+ trials, there were no significant differences in the amplitudes of the late long-latency response across the different times-before-GO conditions. In the SCM-trials, significant differences in the long-latency amplitudes were observed between the GO – 500 and At GO conditions similar to that reported in Figure 3.

**Figure 6:**
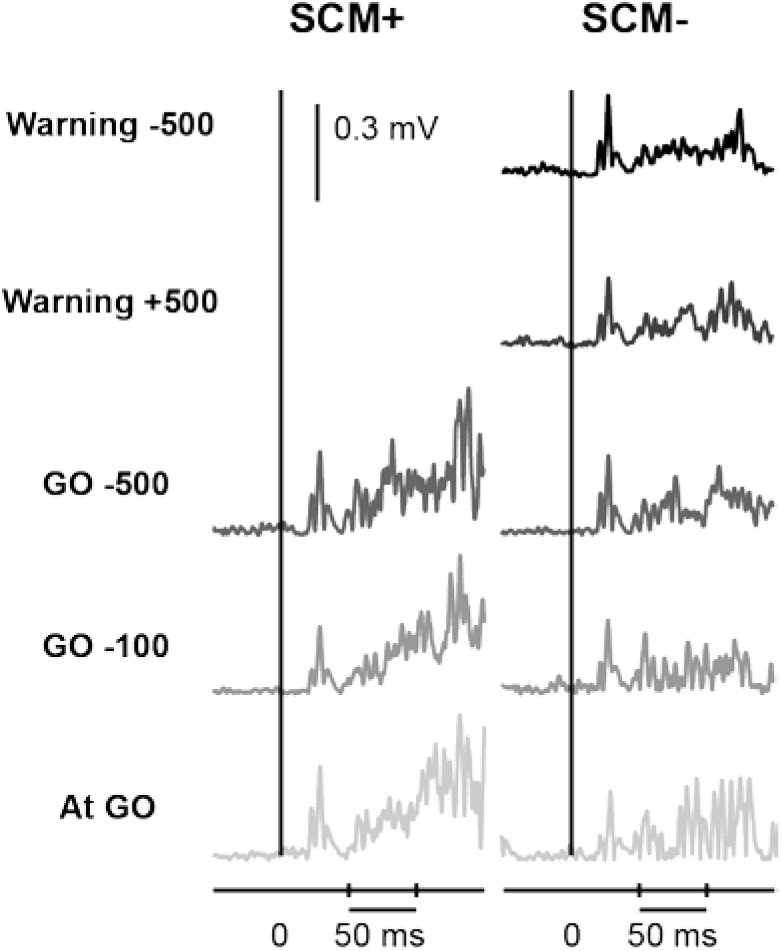
Comparison of triceps EMG response in SCM+ (left) and SCM-(right) trials. Sample data from individual trials in a subject show the response to perturbations applied at different points of time starting at 500 ms before WARNING (top-most) to At GO (bottom-most).

**Figure 7:**
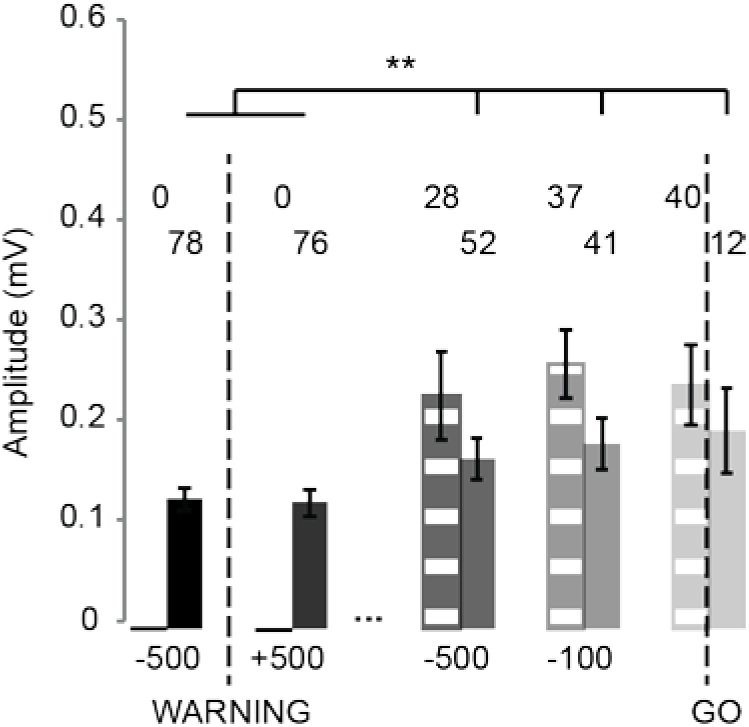
Comparison of the late long-latency response amplitude (group mean ± standard error) of the triceps EMG. Each pair of bars represents the average amplitude of SCM-trials (solid) and SCM+ (hatched). The number of trials included are presented above the columns SCM+ (top) and SCM-(bottom) trials. The time of onset of perturbation is shown along the X-axis from 500 ms before WARNING (left) to perturbation onset At GO (right). The p-values for significant differences are shown.

## Discussion

The purpose of this study was to determine how the nervous system modulates the contributions of LLSR and startReact during the transition from posture to movement. We found that the amplitude of the whole LLSR (without accounting for limb-stabilizing LLSR and startReact components) increases steadily during movement preparation, with no corresponding changes in background activity. The long-latency (particularly late long-latency) response amplitude was most affected during the transition from posture to movement. We found that both the LLSR and startReact were important contributors to the modulation of the LLSR during the transition from posture movement. The LLSR (seen in the early long-latency response and SCM-trials) dominates during the posture phase of movement and is also modulated during the transition to posture. Conversely, the startReact response (SCM+ trials during the late long-latency response) is not a major contributor during the posture phase but becomes increasingly important as movement initiation approaches. Our study distinguishes itself from others that have observed reflex amplitude changes during the transition from maintenance of posture to initiation of movement (Mortimer et al. 1981; Bonnet 1983), by deducing contributions from two separate mechanisms. As these mechanisms are modulated by different structures in the nervous system, this more complete analysis of this complex reflex modulation can provide important insights into deficits following neurological disease and injury.

### Dissociating the LLSR and StartReact

This article distinguishes itself from previous work by accounting for two distinct phenomena: LLSR and startReact. We were the first to demonstrate that the startReact (or triggered reaction) could be identified using the SCM muscle (Ravichandran et al. 2013). This important observation now allows us to evaluate both phenomenon and determine which is dominating the muscular response. The presence of SCM activity indicates the presence of a startReact response. Therefore, modulation observed during SCM-trials can be assumed to be driven by the LLSR. startReact is an early release of a planned movement. It is only evoked when the subject is in a state of motor planning e.g during the transition from posture to movement. Thus, we see no contributions of startReact (i.e. no SCM+ trials) during posture (Figure 7).

It is important that the startReact modulation observed here is not a neural modulation but rather an artifact of averaging. Specifically, SCM+ trials are larger in amplitude that SCM-trials (Figure 7). The SCM+ amplitude does not increase the GO approaches. Rather the likelihood of eliciting a SCM+ increases (Figure 6). The increase observed in the amplitude during the late long-latency period (Figure 3) is the result of averaging more large SCM+ trials with fewer small SCM-trials. Conversely, SCM+ trials could have LLSR represented but the response is dominated by the larger SCM+ response. This result highlights the importance and necessity of evaluating all trials individually and accounts for the long term debate on the role of triggered reactions in the literature because they could not be identified.

### Contributions of LLSR and StartReact during the transition from posture to movement

There is increasing evidence that there are at least two separate components at work during the LLSR (Shemmell et al. 2009). First, a perturbation applied in the presence of a motor plan can trigger an early release of the planned movement (Crago et al., 1976; Gottlieb and Agarwal, 1980; Rothwell et al., 1980). We have previously demonstrated that this effect is similar to the early release of a planned ballistic movement triggered by a startling acoustic stimulus (Ravichandran et al., 2013), the latter which has been studied by numerous groups (Valls-Solé et al., 1999; Siegmund et al., 2001; Carlsen et al., 2004; Honeycutt et al. 2012; Honeycutt et al. 2013; for review, see Valls-Solé et al., 2008). Second, a limb-stabilizing component specific to the mechanics of the task environment (Krutky et al. 2010) and the biomechanics of the limb is present (Kurtzer et al. 2008; Pruszynski et al. 2011). This is the first study to delineate how these separate components are modulated during the transition from posture to movement.

We found that both the LLSR and startReact play key roles in the transition from posture to movement. StartReact appears to dominate the changes immediately prior to movement. The majority of the modulation during the transition phase occurred in the late LLSR, which has significant startReact contributions (Ravichandran et al. 2013). We observed increasing amplitudes in the late LLSR as GO time approached which was tied to the increasing probability of SCM+ trials indicating that this modulation is likely driven by startReact. Importantly as highlighted above, the amplitude of the SCM+ trials during the late long-latency response was not different for the three perturbations time periods before the GO in which SCM+ responses were observed (Fig. 7). Therefore the observed increase in long-latency amplitude with approaching movement onset results from averaging an increasing number of SCM+ trials that are greater in amplitude than SCM-trials.

While startReact dominates changes during the transition from posture to movement, the LLSR was always present and indeed showed modulation during both the early and late long latency time windows. Modulation of the LLSR was also observed in the late long-latency response as the amplitudes of SCM-trials were larger during the movement preparation phase compared to the posture maintenance phase. Still, no differences were found in the SCM-trials during the various time points within the movement preparation phase (Figure 7), indicating that the LLSR alone cannot explain the modulation of the average late long-latency response within this period. Rather the modulation in the late LLSR during the movement preparation phase is likely driven predominately by the presence of startReact.

### Neural mechanisms

The modulation during the LLSR is likely mediated by supraspinal sources and is not a simple spinal gain-scaling. We controlled for background activity in the stretched muscle to rule out gain-scaling. We also observed changes in the short latency response amplitudes only for perturbations delivered coincident with the GO cue. This suggests that the increase in the LLSR we observed for most time periods was not simply due to an increase in background activity or other changes in spinal excitability. Our work is complemented by other studies that controlled for the background activity and also observed no short latency changes in either H-reflexes (Brooke et al. 2000; Hayes and Clarke 1978) and or short-latency stretch reflexes immediately prior to movement (Bonnet 1983; Mortimer et al. 1981; Sullivan and Hayes 1987).

The LLSR and startReact are mediated by distinctive neural structures. There is strong evidence of cortical contributions to the reflex modulation observed when interacting with different mechanical environments (Kimura et al. 2006; Shemmell et al. 2009) and also to compensate for the intersegmental dynamics of the limb (Pruszynski et al. 2011). While the neural origins of startReact has been questioned (Alibiglou and MacKinnon 2012; MacKinnon et al. 2013; Marinovic et al. 2014), current evidence strongly points to a brainstem origin to startReact. Several key papers indicate that the cortex and corticospinal tract are not the primary mediators utilized during startReact. First, individuated movements of the hand that are expressed predominately through the corticospinal tract (Kuypers 1981; Lawrence and Kuypers 1968; Lemon et al. 2012; Schieber 2011; 2004) are not susceptible to early release by startle (Carlsen et al. 2009; Honeycutt et al. 2013). Second, startReact is intact in patient populations with cortical (stroke survivors) and corticospinal (hereditary spastic paraplegic patients) damage (Honeycutt and Perreault 2012; Nonnekes et al. 2014) showcasing that degradation to these pathways does not eliminate the early release of movement via startle.

Based on that evidence and similarities between the acoustic stimulus-evoked and perturbation-evoked early onsets of motor plan, we suggest that the brainstem is the most likely origin for perturbation-evoked early onsets that accompany muscle activity in the SCM muscle. Consistent with this, both preparatory and movement related activity is observed in the reticular formation of the brainstem of cats (Schepens and Drew 2004) and monkeys (Buford and Davidson 2004). This is further strengthened by the observation of activity in the reticular formation immediately preceding a startle-like response triggered by removal of the limb support surface in standing cats (Stapley and Drew 2009). There is also evidence for excitatory connections between the giant neurons in the reticular formation and the limb motor neurons in animals (Lingenhohl and Friauf 1994; Yeomans and Frankland 1995), which could provide the necessary pathways linking the brainstem with the early release of voluntary movement.

It has been well documented that the motor cortex can also respond to external perturbations of posture and that its response depends on the prepared motor state (Evarts and Tanji 1976; Pruszynski et al. 2014; Tanji and Evarts 1976). Indeed, tasks such as individuated finger movements, thought to be mediated by the motor cortex, can exhibit responses that behave similar to triggered reactions except acting more slowly and unrelated to the presence or absence of simultaneous brainstem activity (Carlsen et al. 2009; Honeycutt et al. 2013). How these cortical pathways work together with brainstem pathways for the many tasks in which both can contribute remains an interesting topic for future research.

### StartReact in other paradigms

It is important to note that our observations about the importance of startReact during the long-latency stretch reflex time window may not translate to other paradigms. For example, recent work from Foggard et. al. demonstrates long-latency stretch reflex modulation but poor or no influences of startle/startReact (Castellote et al., 2017; Forgaard et al., 2015; Forgaard et al., 2016). This work, like our own, highlights that the long latency stretch reflex window can be modulated in the absence of startReact. Furthermore, it highlights how differences in paradigms can relate to big differences in the role of startReact. For example, only unexpected perturbations elicit startle (Forgaard et. al. 2016). However, perhaps the most important contribution of this work is that in “neck activity” can be elicited as part of a postural response to the perturbation itself (Forgaard et. al. 2016). Thus, all future work that attempts to evaluate the role of the startReact reflex on reflex contributions must necessarily ensure that the responses of the neck are true “SCM+” responses and not the result of postural control. Unlike Forgaard’s work, we rarely (<2% of trials) see activity in the neck during voluntary or “do not intervene” conditions. Therefore, neck activity is only present when a perturbation is presented in the presence of a movement plan. Therefore, our particular paradigm appears to have less postural influence. Still, those who wish to further evaluate the influences of startReact should carefully consider Forgaard’s work and the influence of “neck” postural reflexes.

While the contradictions or differences between paradigms could be perceived as negatives, we would argue that they allow for great opportunity. As we have pointed out, a positive startReact does not preclude other forms of modulation but simply that startReact is also influencing the response. Therefore, having different paradigms that elicit strong startReact (our paradigm) and those that have minimal or no influence of startReact (Forgaard et. al. 2016), will be critical as we work to fully understand the role of the underlying mechanisms contributing to reflexive modulation.

## Conclusions and Functional Implications

Our study showcases that the increase in the EMG amplitude observed prior to a voluntary movement is due to contributions from both the LLSR and startReact working together to generate a sophisticated response. Modulation in the early long-latency response is likely dominated by the LLSR while the observed increase in the late-long latency response is dominated by startReact. Having a clear understanding of which components contribute during this response is most important in light of patient populations that have lesions affecting different parts of the nervous system. For example, stroke patients are likely to have more significantly impaired LLSRs, given their cortical origin. In conclusion, the present findings highlight the need to consider the multiple pathways that can contribute to reflex responses in the long-latency time period, and emphasize the importance of interpreting these reflexes and their associated pathways in a task-specific manner.

## Acknowledgments

The authors would like to thank Tim Goetz-Haswell his technical and scientific expertise. This work was supported by the National Institutes of Health grants R01 NS053813 and R00 HD073240.

